# Identification of a fully dechlorinated product of chlordecone in soil microcosms and enrichment cultures

**DOI:** 10.1101/2021.03.03.433826

**Authors:** Line Lomheim, Robert Flick, Suly Rambinaising, Sarra Gaspard, Elizabeth A. Edwards

## Abstract

Anaerobic microcosms constructed with soil from Guadeloupe, amended with electron donor (ethanol and acetone) and incubated for more than a decade, transformed chlordecone (CLD) into a suite of progressively more dechlorinated products, including a fully dechlorinated carboxylated indene product. This fully dechlorinated transformation product has never before been observed and indicates that complete dechlorination of CLD is possible. The carboxylated indene was detected by LC-MS and structure was confirmed by LC-MS/MS using a Q-Exactive Orbitrap mass spectrometer.

## INTRODUCTION

Chlordecone (C_10_Cl_10_O) is an organochlorine pesticide first synthesized in 1951 and commercially introduced to the United States in 1958 by Allied Chemical under the name Kepone®. From 1951 to 1975, U.S. production of chlordecone was 1,600 tons. Use in the U.S. was banned in 1974, but continued elsewhere. Chlordecone use continued in Guadeloupe and Martinique between 1971 and 1993 to control the banana weevil, *Cosmopolitans sordidus*, and the amount of chlordecone applied over this time is estimated to be 300 tonnes.^1^ As a result of long term use and very slow degradation rates, plantation soils, surface and groundwater runoff, and coastal marine waters are still polluted with chlordecone.^2, 3^ Plants, land and marine animals in contact with polluted soils and waters are also impacted^4, 5^ and consumption of contaminated food may not be without health consequences for local populations. Chlordecone is carcinogenic and has endocrine disrupting properties.^6, 7^ Over 90% of the population in Guadeloupe and Martinique have detectable CLD in their blood.^6, 7^

Soil, groundwater and surface contamination in Martinique and Guadeloupe has been delineated to track the fate of chlordecone.^2^ Activated charcoal filtration has been implemented to reduce exposure from drinking water.^8^ However, many soils remain contaminated and although efforts are underway to find ways to remove chlordecone from polluted soils, there is little success. Sequestration, phytoremediation and bioremediation options have been investigated.^4, 9–13^, Current approaches to minimize health impacts involve modifications to agricultural practices, such as avoiding pasturing animals on contaminated soils and restricting what can be grown or harvested for consumption.^3^

Recent studies are providing some hope that agricultural soils could be managed through stimulating reductive dechlorination of chlordecone. Reductive dechlorination of chlordecone using vitamin B12 has been known since 1978.^14^ In the laboratory, reductive conditions can be established abiotically by adding reduced vitamin B12,^8^ zero valent iron,^15^ or biologically through addition of electron donor to anaerobic sulfate-reducing or methanogenic enrichment cultures or isolates.^10, 11, 16–19^ Products formed during reductive dechlorination in laboratory studies have also been detected in samples from polluted agricultural soils,^10, 16^ suggesting that some natural anaerobic transformation is occurring *in situ*, albeit very slowly. The anaerobic transformation products observed to-date include mono-, di-, and tri-hydrochlordecone, polychloroindenes as well as carboxylated polychlorinated indenes (Figure 1). These families of transformation products are very well detailed in recent publications.^10, 11, 18^ We developed an LC/MS analysis method needing only small volumes and simple sample preparation protocols that was applied to a series of anaerobic soil microcosms we had set up years earlier.^16^ Nineteen different chlordecone transformation products were detected in biologically active microcosms, including hydrochlordecones, and open-cage polychlorinated indenes and polychlorinated carboxylated indenes. To our surprise we detected products corresponding to removal of up to 9 chlorine atoms.^16^ In a recent study using a pure culture of *Desulfovibrio*, Della-Negra *et al.,*^17^ identified these same families of transformation products plus additional thiol derivatives generated by bacterial reductive sulfidation in the presence of reduced forms of sulfur. These thiol derivatives were also detected in samples from chlordecone-contaminated mangrove bed sediments in Martinique.^17^ The biochemical pathways and mechanisms leading to the formation of all these various families of biological transformation products have not yet been established.

**Figure 1.**
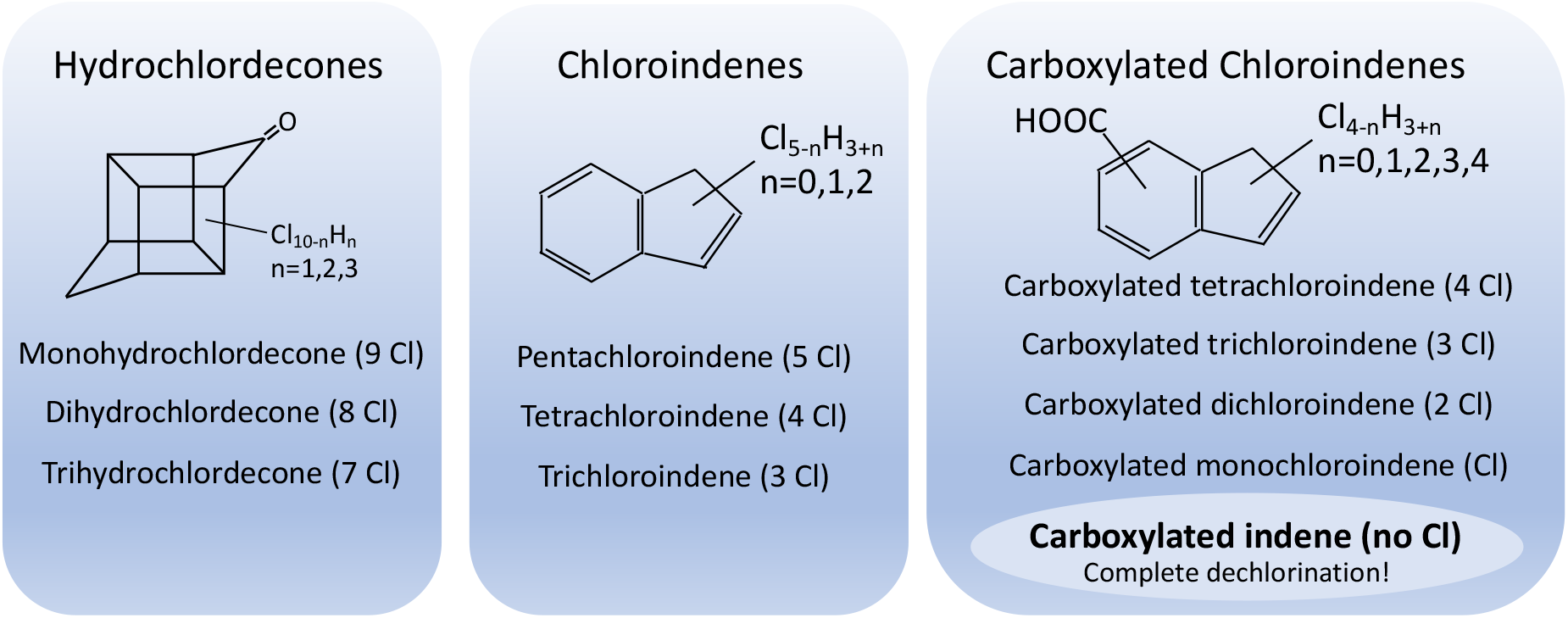
Three groups of dechlorination products observed in Guadeloupe microcosms: hydrochlordecones (A-compounds, bishomocubane structure); chloroindenes (B-compounds, opened ring structure); and carboxylated chloroindenes (C-compounds, opened ring structure with carboxyl group).

In this short communication, we report on the continued investigation of the biotransformation of chlordecone in a series of anaerobic microcosms and transfer cultures that have been incubated for over a decade.^16^ LC/MS analyses had previously revealed extensive dechlorination, and in particular a compound with a mass to charge ratio (m/z) that suggested a fully dechlorinated carboxylated indene product. In this study we used a modified extraction method to analyze new samples from these same microcosms and transfer cultures using tandem LC-MS/MS. Fragmentation spectra were compared to those of available authentic standards to provide unequivocal evidence for production of completely dechlorinated products. Fully dechlorinated carboxylated indene transformation products have never before been observed and suggest that complete dechlorination of CLD is possible. The LC-MS/MS analyses also further supported the proposed structures of all chlorinated transformation products previously detected in these microcosms, transfer cultures and Guadeloupe soil samples.

## MATERIALS AND METHODS

### Chemicals

CLD (neat) and CLD standard (analytic standard 1mg/mL in methanol) were purchased from Accustandar (New Haven, USA). A carboxylated indene standard (1H-Indene-3-carboxylic acid) was purchased from Sigma-Aldrich (catalog #MNO000013). Hexanes and acetone (Fisher), water and methanol (Caledon Laboratory Chemicals), were all of HPLC grade. A pentachloroindene standard (2,4,5,6,7-pentachloro-1H-indene) referred to as B1, was kindly provided by Dr. Pierre-Loïc Saaidi at Génoscope (Évry, France). Chemicals and samples were manipulated with syringes, pipettes, centrifuge tubes and vials all made of glass to minimize any sorption.

### Microcosms and transfer cultures

Soil and water samples used to prepare microcosms were collected from several locations near a banana plantation in Guadeloupe in the fall of 2010. Original microcosms were set up in late 2010 in 160 mL glass serum bottles sealed with blue rubber stoppers while later transfers made into 250 ml glass bottles sealed with screw-cap Mininert™ valves (Chromatographic Specialties). The two active microcosms (G4 and G19), five transfer cultures (GT3, GT4, GT5, GT20, GT33) and four mercuric chloride-poisoned control microcosms (G28, G30, G41, G50) described in this study were part of a larger published experiment;^16^ microcosm names are consistent with this previous publication. The microcosms were amended with CLD at a target nominal concentration of 10 mg/l. Bottle GT3 was also amended with TCE at 10 mg/l as a positive control for reductive dechlorination. All bottles received an electron donor mix of acetone and ethanol, repeated as necessary. The microcosms have been incubated in the dark, unshaken in an anaerobic glovebox since 2010. In 2013, a subset of active microcosms were transferred into larger bottles and topped up with anaerobic mineral medium. Since 2013, original microcosms (G4 and G19) were no longer amended with donor or chlordecone, but were still incubated under anaerobic conditions. The five microcosm transfers (GT3, GT4, GT5, GT20, GT33) continued to be re-amended with donor and acceptor regularly. Additional details regarding setup, amendments and monitoring can be found in our previous publication.^16^.

### Sampling, extraction and LC-MS and LC-MS/MS analyses

Shaken samples (5 ml) containing a slurry of soil and water were sampled in October 2019 and September 2020 from designated bottles. These samples were acidified (pH~1.5) with HCl before they were extracted two times using 5 ml of a mix of 15% (by vol) acetone and 85% hexane. The extracts were evaporated to dryness under a stream of nitrogen and the samples were re-dissolved in 0.5 ml methanol and filtered through a 0.2 um Millex PTFE syringe filter (Millipore Sigma) into 2 ml glass autosampler vials with open top screw cap and PTFE lined septum (Chromatographic Specialties). Chromatography was carried out on a ZORBAX Rapid Resolution High Definition Phenyl-Hexyl column (50mm x 3.0mm, 1.8um) (Agilent, Santa Clara, USA) equipped with a guard column, using a Thermo Scientific Ultimate 3000 UHPLC (Thermo Fisher Scientific, Waltham, MA). The column temperature was 40°C and the flow rate was set to 300 μL^∂^min^−1^. The eluents used were water (A) and methanol (B), and both eluents contained 5 mM of ammonium acetate. The gradient started at 50% B, followed by a linear gradient to 100% B over 8 min, then a hold at 100% B for 4 min, then a return to 50% B over 1 min, and finally a re-equilibration under the initial conditions of 50% B for 5 min (total runtime 18 min). Liquid samples (10 μL) were injected using an Ultimate 3000 UHPLC autosampler, with autosampler temperature of 8°C. Compounds were detected and quantified using a Q-Exactive Orbitrap mass spectrometer (Thermo Fisher Scientific) equipped with a Heated Electrospray Ionization (HESI II) probe, operating in negative ionization mode. Full MS spectra were acquired over an m/z range from 100 to 750 with the mass resolution set to 70k, and common setting parameters were as follows; AGC Target: 3E6, max injection time 200 ms, spray voltage 3.5 kV, capillary temperature 320°C, sheath gas 15, aux gas 5, spare gas 2, and s-lens RF level 50. Parallel Reaction Monitoring (PRM) of the compounds (see Table S1 for details) was carried out at a resolution of 17.5k, and common setting parameters were as follows: AGC Target 2e5, max injection time 100ms, isolation window of 0.5 m/z, with Higher energy Collisional Dissociation (HCD) performed at 35% normalized collision energy (NCE). Data were processed through Xcalibur Qual Browser (Thermo Fisher Scientific), with fragmentation patterns compared to reference standards for each class of compounds. Calibration standards of CLD, pentachloroindene B1 (2,4,5,6,7-pentachloro-1H-indene) and carboxylated indene (1H-Indene-3-carboxylic acid) were prepared from successive dilutions in a 50/50% mix of methanol and mineral medium.

## RESULTS AND DISCUSSION

### Identification of a fully dechlorinated product by LC-MS/MS

Using the modified extraction protocol described above, we successfully detected the same three families of transformation products as previously reported. These are the hydrochlordecones (A-compounds), chloroindenes (B-compounds) and carboxylated chloroindenes (C-compounds) illustrated in Figure 1. Retention times, mass-to-charge (m/z) ratios, and molecular formulae are provided in Table S2.

As previously reported, an extended suite of progressively more dechlorinated products were identified in all seven microbially-active anaerobic bottles, but not in the four poisoned control microcosms (Table S3). In addition to full scan data, we further refined proposed compound structures using LC-MS/MS. We secured authentic standards for chlordecone, one isomer of pentachloroindene and one isomer of a carboxylated indene. The retention time, mass spectra and fragmentation patterns for each standard were compared to the products found in samples. In this way, we were able to identify a fully dechlorinated CLD product: a carboxylated indene (C_10_O_2_H_8_) (Figure 2). The full scan and fragmentation patterns are identical to the 4^th^ decimal place. The retention time of the peak in the sample (1.3 mins) is slightly longer than that of the standard (1.2 mins), suggesting that the product in our samples is not the same isomer as the available standard. This non-chlorinated carboxylated chloroindene was detected in five of the seven microbially-active bottles (Table S3). Under the conditions imposed in these microcosms, we have found for the first time that CLD is slowly transformed by the action of an indigenous anaerobic microbial community to a fully dechlorinated product.

**Figure 2.**
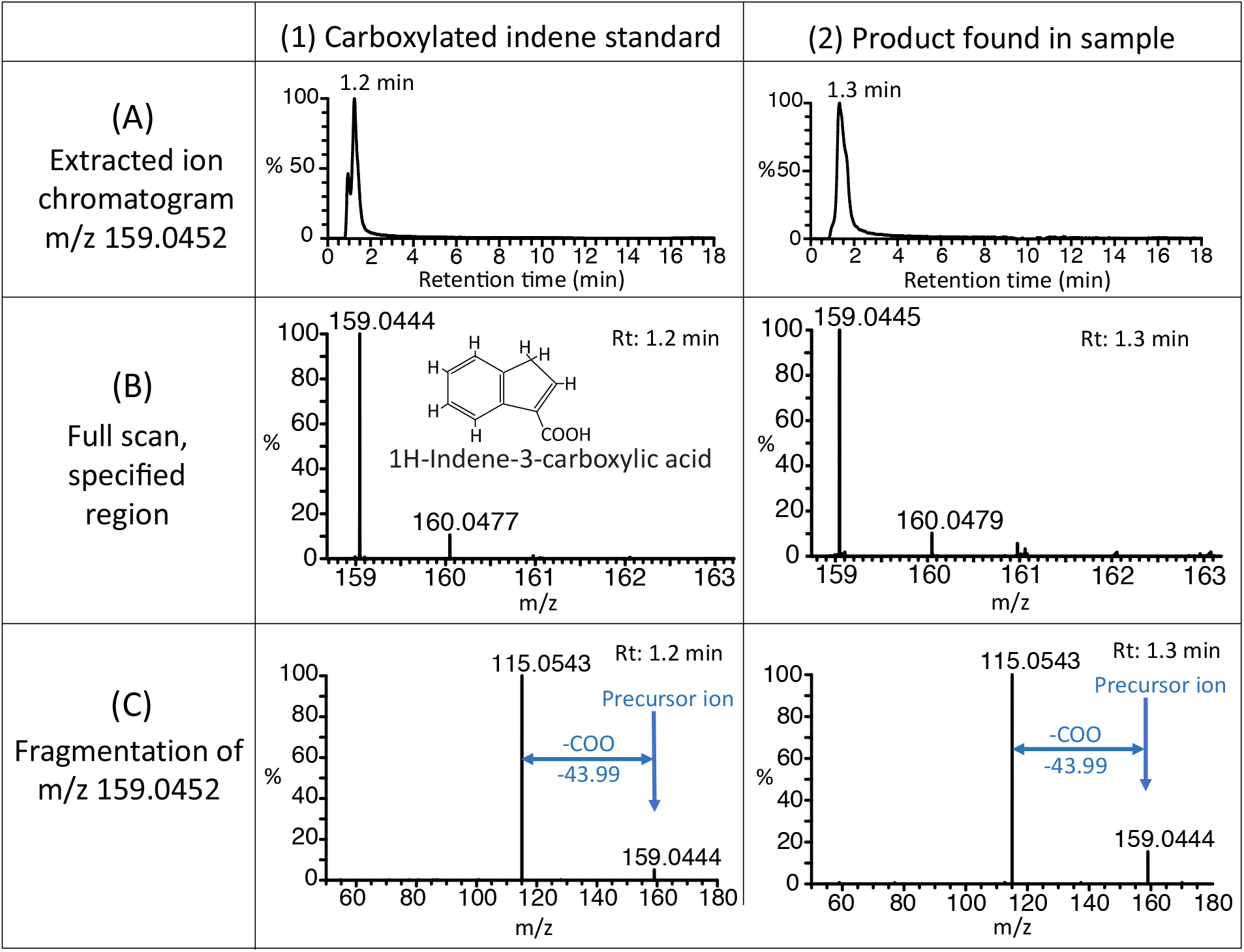
Identification of carboxylated indene [C_10_O_2_H_7_]^−^ by LC-MS and LC-MS/MS in microcosm constructed with soil from Guadeloupe. (A) extracted ion chromatograms m/z 159.0452, (B) LC-MS mass spectra (full scan, region m/z 159-163), and (C) LC-MS/MS mass spectra from the fragmentation of m/z 159.0452, for (1) a carboxylated indene standard, Rt 1.2 min and (2) a microcosm sample (G19), Rt 1.3 min, all run in negative mode.

### Structural elucidation of partially dechlorinated and decarboxylated products by LC-MS/MS

LC-MS/MS analysis also provided support for the structures of previously observed parent molecules and partially dechlorinated products. LC-MS and MS/MS spectra of a chlordecone standard compared to peaks assigned as chlordecone and monohydrochlordecone (H replacement of one Cl) and dihydrochlordecone (replacement of two Cl substituents) products observed in Guadeloupe microcosm and transfer culture samples are shown in Figure S1. The fragmentation spectra all show loss of m/z of 79.97 from their parent ion, corresponding to COOHCl.

The carboxylated chloroindenes are an interesting aromatic ring-opening product that maintains all 10 carbons from the parent chlordecone molecule but with a carboxyl substituent presumably derived from the orginal ketone, and much lower number of chlorine substituents. Multiple isomers are possible (Table S2). LC-MS and LC-MS/MS spectra of mono, di-, tri- and tetra-chlorinated indene carboxylates identified in Guadeloupe microcosm samples are shown in Figure S2. The spectra for the newly discovered completely dechlorinated carboxylated indene product and standard presented in Figure 2, are included again in Figure S2 to provide comparison to other spectra.

Chlordecone or partially dechlorinated products must also undergo some kind of decarboxylating reaction to produce 9 carbon chloroindenes. LC-MS and MS/MS spectra of a pentachloroindene standard compared to three peaks assigned to chlorinated indene products with 5, 4 and 3 chlorine substituents observed in Guadeloupe microcosms are shown in Figure S3. The concordance between standard and sample supports the identity of each of the classes of transformation products, although the existence of multiple structural isomers means that the position of substituents around the ring cannot be determined.

As mentioned previously, multiple isomers of certain metabolites were previously identified, having exactly the same m/z ratio but varying retention time. By examining the LC-MS/MS mass spectra of these isomeric compounds, we were able to verify that their LC-MS/MS spectra were indeed the same (Figure S4). We can therefore conclude with even more confidence that our previous isomeric metabolite assignments were correct.

### Transformation product concentration estimations

We used the three standards available to generate calibration response factors (Table S4) to then use to estimate concentrations (Table S3b). The chlordecone standard was used to quantify chlordecone and hydrochlordecones, the pentachloroindene standard was used for all chlorinated indenes, and carboxylated indene standard was used for all chlorinated and non-chlorinated carboxylated indenes. We then estimated overall recovery as the sum of the moles of all measured products plus remaining CLD at the time of sampling (Oct. 2019) divided by the total moles CLD added to the bottle since 2010. This overall recovery over ~9 years ranged from 19-82% (Table S3b), which seems acceptable given the length of time elapsed. These calculations also show that measured products accounted for 9% to 67% of the total added chlordecone, depending on the bottle. We did not account for mass removed at each sampling time. The molar proportions of each of the compounds analyzed in each microcosm bottle are depicted in Figure 3 (Bottles G19 and GT20) and in Figures S5a-e for remaining bottles. Patterns are very similar in all active bottles but not in poisoned controls where only chlordecone (>99%) and small concentrations of hydrochlordecone were detected (Table S3). In our previous study, aqueous samples or slurries were simply mixed with methanol, without concentration.^16^ Samples in this study were extracted with acetone/hexane and then concentrated by evaporation. This was necessary to obtain LC-MS/MS fragmentation patterns. Although we achieved more mass on-column, the new extraction method had about 2 times lower overall recovery efficiency for CLD, hydrochlordecone and carboxylated indenes, and 5 times better recovery of pentachlorindene compared to our previously published data (Table S5).

**Figure 3.**
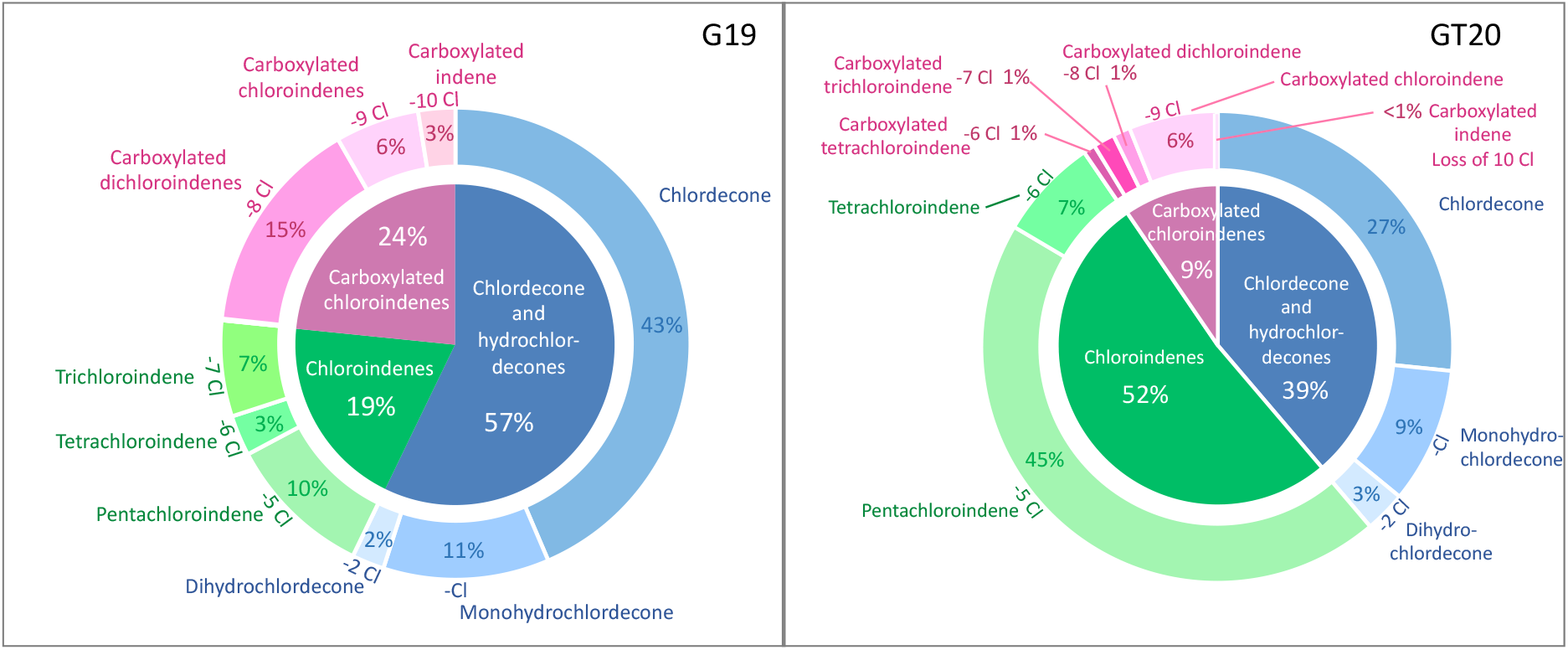
Estimated molar concentration distribution of chlordecone and dechlorinated metabolites in one original microcosm (G19; left) and in a transfer culture (GT20; right) after a decade of anaerobic incubation. Charts were constructed based on measured and estimated concentrations in nmol/l (Table S3b).

### Broader implications

These results illustrate the potential for reductive transformation of chlordecone into less chlorinated aromatic fused ring intermediates and at least one fully dechlorinated aromatic product. While an overall mass balance does not suggest that complete mineralization is occurring under these strictly anoxic conditions, the process does offer some hope that partial dechlorination and transformation could result in products that are more amenable to aerobic biodegradation. Indeed, aromatic compounds like indene can be metabolized aerobically^20, 21^ through the action of various oxygenases.

The pathways leading to the formation of the observed metabolites, as well as the mechanisms for these biotransformations are not known. Reductive dehalogenases and organohalide respiring bacteria have not been implicated, although the involvement of corrinoids has been proven in one case.^22^ The product distribution maps shown in Figure 3 are both intriguing and confusing. Intriguing because following clockwise around the circle one finds progressively fewer chlorine substituents, suggesting a stepwise process. However, the chlorinated indenes (9C) are more chlorinated than the carboxylated indenes (10C), which is confusing if the carboxyl substituent remained from the parent compound. Perhaps, multiple separate pathways to the various sets of products are operating, unless carboxylation occurs following formation of pentachloroindene.

Finally, the rates of transformation in these particular microcosms are quite slow. Other researchers have found considerably faster rates in much more enriched or purified cultures.^10, 17, 22, 23^ The stage is set now to determine first if these products are less toxic and problematic than chlordecone itself, and next how to reproduce such strictly anoxic conditions for treatment of agricultural soils. Perhaps through intermittent flooding and donor addition, reductive dechlorinating transformation can be accelerated. These metabolites should be monitored in the field to establish the extent of natural transformation.

## ASSOCIATED CONTENT

### Supporting Information

Supporting Method Details (SMD) for LC-MS/MS method. Figures showing LC-MS and MS/MS ion spectra for compounds belonging to each of the three metabolite groups, LC-MS and MS/MS ion spectra for isomers of carboxylated chloroindenes, and pie charts of estimated distribution of chlordecone and dechlorinated metabolites in microcosm samples (microcosm samples that were not presented in Figure 3). Tables showing additional LC-MS/MS method information, details of chlordecone and dechlorinated metabolites identified in microcosm samples, raw data, estimated concentrations and calibration curves from LC-MS analysis, and comparisons of results from two different sample preparation methods.

## ACKNOWLEDGEMENTS

Funding was provided by a grant from the Préfecture de Martinique, under the auspices of plan Clordécone III, ACTION « remediation » (2019 PITE BOP 162) and the Natural Science and Engineering Research Council (NSERC) of Canada.

## Notes

### Competing Interest Statement

The authors have declared no competing interest.

