## Supplemental information for "Identification of a fully dechlorinated product of chlordecone in soil microcosms and enrichment cultures"

**TABLE OF CONTENTS**

**LIST OF SUPPORTING FIGURES**

**Figure S1:** LC-MS and MS/MS spectra of a chlordecone standard and of chlordecone and two hydrochlordecone products observed in Guadeloupe microcosms. Examples are from microcosm G19.

**Figure S2:** LC-MS and LC-MS/MS spectra of a carboxylated indene standard and of three carboxylated chloroindene/carboxylated indene products observed in Guadeloupe microcosms. Examples are from microcosms G19 (carboxylated indene, carboxylated chloroindene and carboxylated dichloroindene) and GT20 (carboxylated trichloroindene and carboxylated tetrachloroindene).

**Figure S3:** LC-MS and MS/MS spectra of a pentachloroindene standard and of three chloroindene. products observed in Guadeloupe microcosms. Examples from microcosm G19.

**Figure S4:** Examples of chromatograms and mass spectra from LC-MS/MS runs for four metabolites in the carboxylated chloroindene family for which we have observed multiple isomers; carboxylated tetrachloroindene, carboxylated trichloroindene, carboxylated dichloroindene and carboxylated chloroindene.

**S4a**: Comparison of LC-MS/MS ion spectra of the fragmentation of m/z 294.8893 (carboxylated tetrachloroindene) at $t_R$ 4.62 min and 5.03 min (sample GT20).

**S4b**: Comparison of LC-MS/MS ion spectra of the fragmentation of m/z 260.9282 (carboxylated trichloroindene) at $t_R$ 2.99 min, 3.47 min, 5.83 min and 6.38 min (sample GT20).

**S4c**: Comparison of LC-MS/MS ion spectra of the fragmentation of m/z 226.9672 (carboxylated dichloroindene) at $t_R$ 3.76 min, 4.31 min and 4.97 min (sample G19).

**S4d**: Comparison of LC-MS/MS ion spectra of the fragmentation of m/z 193.0062 (carboxylated chloroindene) at $t_R$ 2.43 min and 2.81 min (sample G19).

**Figure S5 a-e:** Estimated distribution of chlordecone and dechlorinated metabolites in microcosms and transfer cultures (a) G4, (b) GT3, (c) GT5, (d) GT33 and (e) GT4. Charts were constructed from measured and estimated concentrations in nmol/l (see Table S3b for details).

### LIST OF SUPPORTING TABLES

*All Supporting information tables are provided in spreadsheet format in the accompanying Excel file*

**Table S1:** LC-MS/MS Parallel Reaction Monitoring (PRM) compound table (parameters used in LC-MS/MS run)

**Table S2:** Chlordecone and dechlorinated metabolites identified in microcosm and enrichment samples

**Table S3 a-c:** LC-MS results; (a): Raw data (area count), (b): Estimated concentration (nmol/l of slurry) and (c): Standards run and response factors used

**Table S4 a-d:** Standards and Calibration; Calibration curves for (a) chlordecone, (b) pentachloroindene (2,4,5,6,7-pentachloro-1H-indene), and (c) carboxylated indene (1H-Indene-3-carboxylic acid), and (c) raw data for all standards

**Table S5 a-**b: Comparing results from two different sample preparation methods (for comparison to previously published data)

### SUPPORTING FIGURES

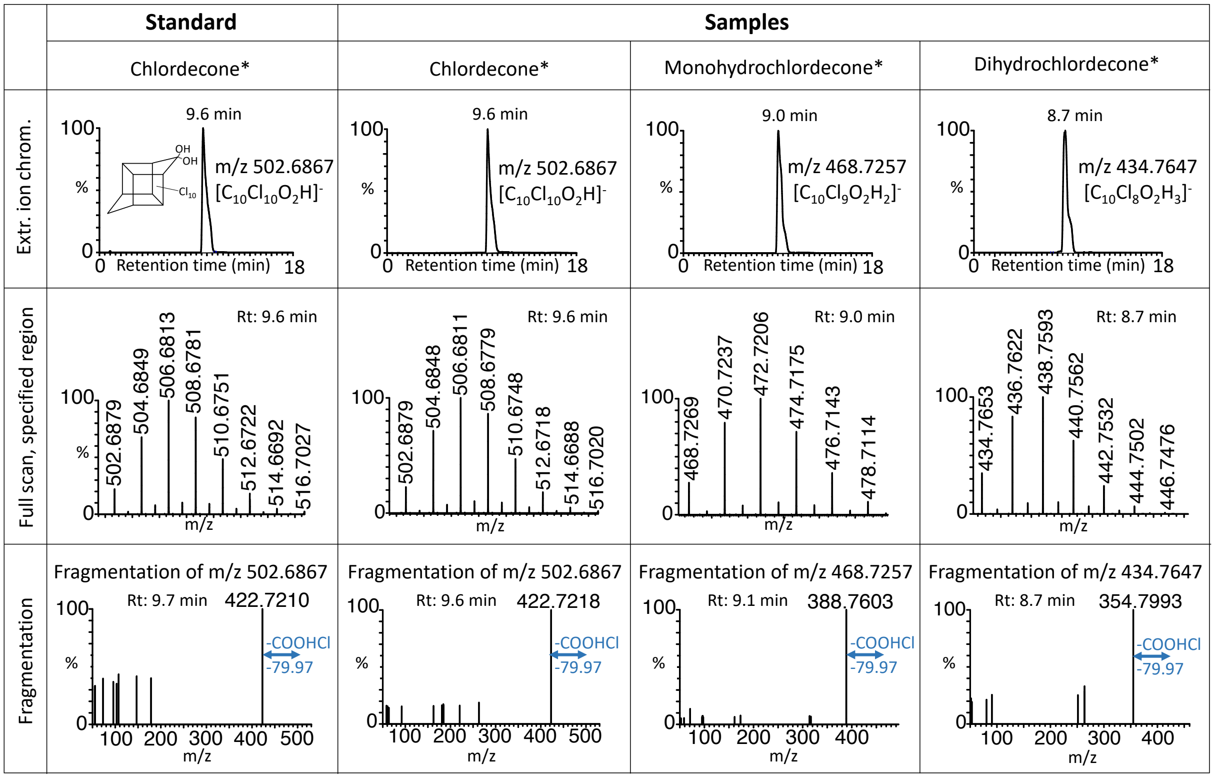


* Hydrate form of chlordecone, monohydrochlordecone and dihydrochlordecone

**Figure S1: LC-MS and MS/MS spectra of a chlordecone standard and of chlordecone and two hydrochlordecone products observed in Guadeloupe microcosms.** Mass spectra were obtained from the fragmentation of m/z 502.6867 in a chlordecone standard (Rt=9.6 min) and m/z 502.6867 chlordecone $[C_{10}Cl_{10}O_2H]^-$ (Rt=9.60 min), m/z 468.7257 monohydrochlordecone $[C_{10}Cl_9O_2H_2]^-$ (Rt=9.10 min), m/z 434.7647 dihydrochlordecone $[C_{10}Cl_8O_2H_3]^-$ (t=8.74 min) in microcosm samples. Examples are from microcosm G19.

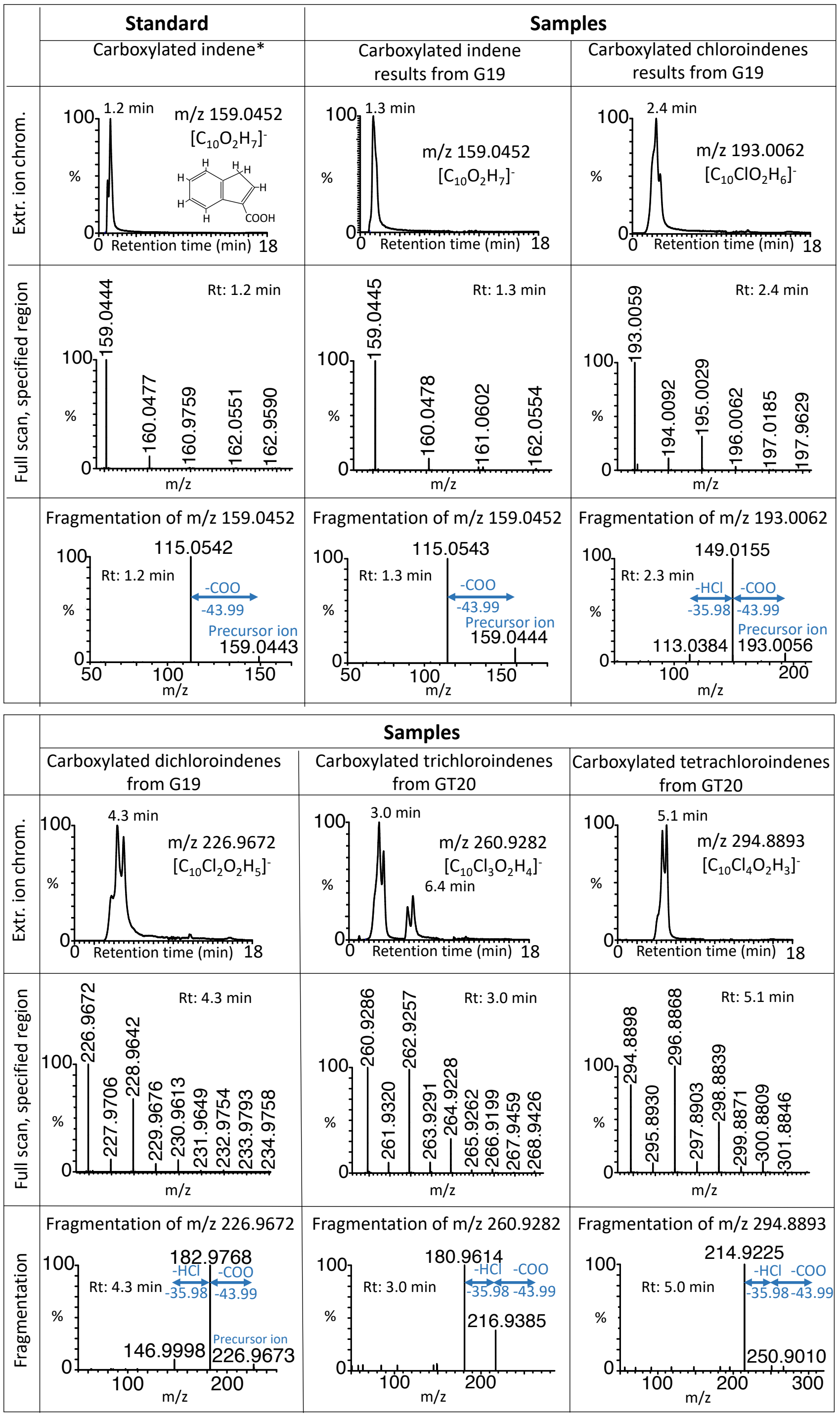


* 1H-Indene-3-carboxylic acid

**Figure S2: LC-MS and LC-MS/MS spectra of a carboxylated indene standard and of three carboxylated chloroindene/carboxylated indene products observed in Guadeloupe microcosms**. Mass spectra were obtained from the fragmentation of m/z 159.0452 in a carboxylated indene standard (t=1.26 min) and m/z 159.0452 carboxylated indene $[C_{10}O_2H_7]^-$ (t=1.3 min), m/z 193.0062 carboxylated chloroindene $[C_{10}ClO_2H_6]^-$ (t=2.4 min), m/z 226.9672 carboxylated dichloroindene [$C_{10}Cl_2O_2H_5$]$^-$ (t=4.67 min), m/z 260.9282 carboxylated trichloroindene [$C_{10}Cl_3O_2H_4$]$^-$ (t=2.99 min), m/z 294.8893 carboxylated tetrachloroindene [$C_{10}Cl_4O_2H_3$]$^-$ (t=4.98 min) in microcosm samples. Examples are from microcosms G19 (carboxylated indene, carboxylated chloroindene and carboxylated dichloroindene) and transfer GT20 (carboxylated trichloroindene and carboxylated tetrachloroindene).

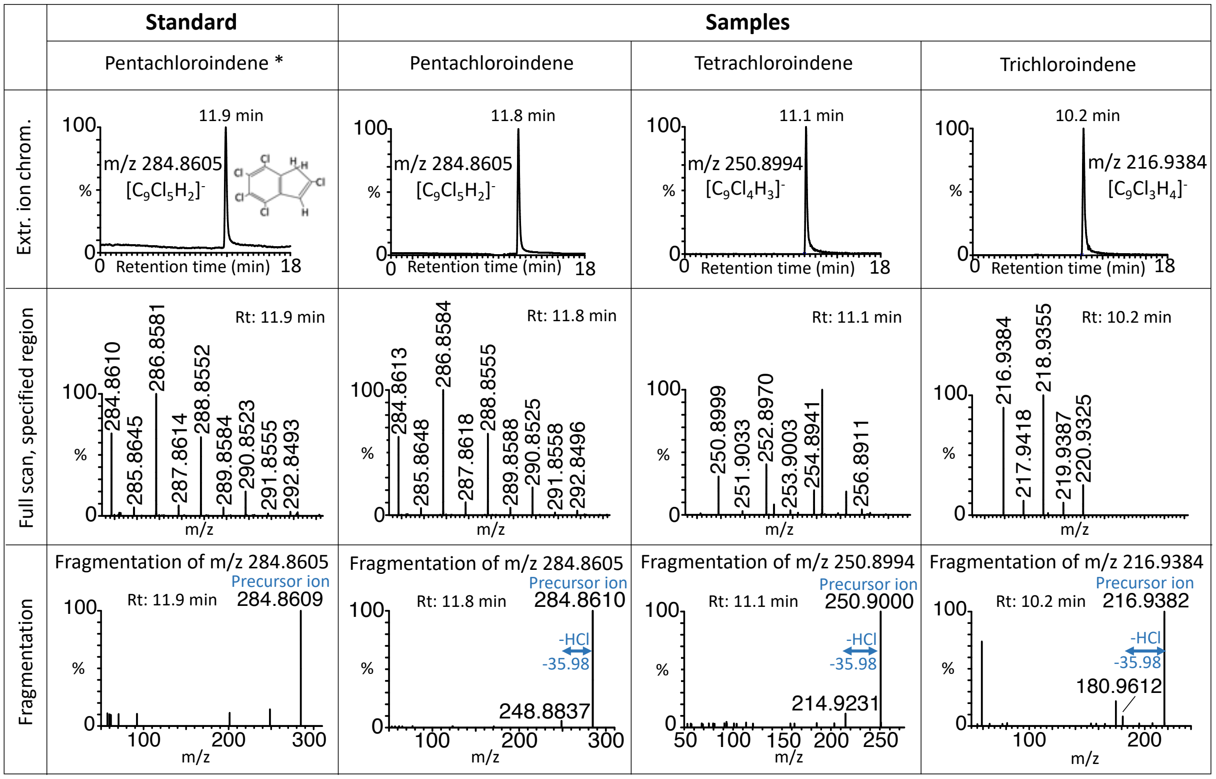


* 2,4,5,6,7-pentachloro-1H-indene

**Figure S3: LC-MS and MS/MS spectra of a pentachloroindene standard and of three chloroindene products observed in Guadeloupe microcosms**. Mass spectra were obtained from the fragmentation of m/z 284.8605 in a pentachloroindene standard (Rt=11.93 min) and m/z 284.8605 pentachloroindene [$C_9Cl_5H_2$]$^-$ (Rt=11.83 min), m/z 250.8994 tetrachloroindene [$C_9Cl_4H_3$]$^-$ (Rt=11.16 min), m/z 216.9384 trichloroindene [$C_9Cl_3H_4$]$^-$ (Rt=10.28 min) in microcosm samples. Examples are from microcosm G19.

**Explanation for Figure 4a-d: Verifying the presence of isomers in the carboxylated indene family by comparing their LC-MS/MS spectra.**

Multiple isomers of certain metabolites were previously identified, having exactly the same mass to charge ratio but varying retention time. By examining the LC-MS/MS mass spectra of these isomeric compounds, and verifying that their LC-MS/MS spectra are the same, we were able to increase our confidence that our previous isomeric metabolite assignments are correct.

**Figures S4a-d** show examples of chromatograms and mass spectra from LC-MS/MS runs for four metabolites in the carboxylated chloroindene family for which we have observed multiple isomers; carboxylated tetrachloroindene, carboxylated trichloroindene, carboxylated dichloroindene and carboxylated chloroindene. Examples in are from microcosm GT20 for carboxylated tetrachoroindene (two distinct peaks) and carboxylated trichloroindene (three distinct peaks), and from G19 for carboxylated dichloroindene (three distinct peaks) and carboxylated chloroindene (two distinct peaks).

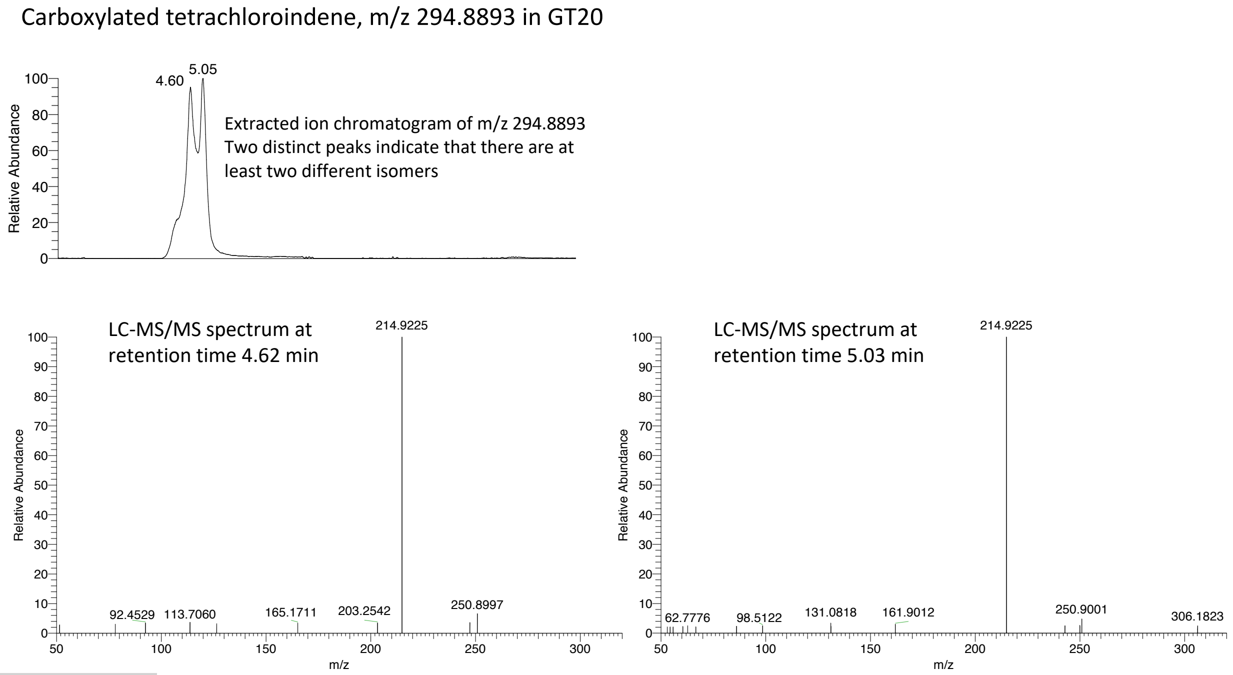


**Figure S4a**: Comparison of LC-MS/MS ion spectra of the fragmentation of m/z 294.8893 (carboxylated tetrachloroindene) at $t_R$ 4.62 min and 5.03 min (sample GT20).

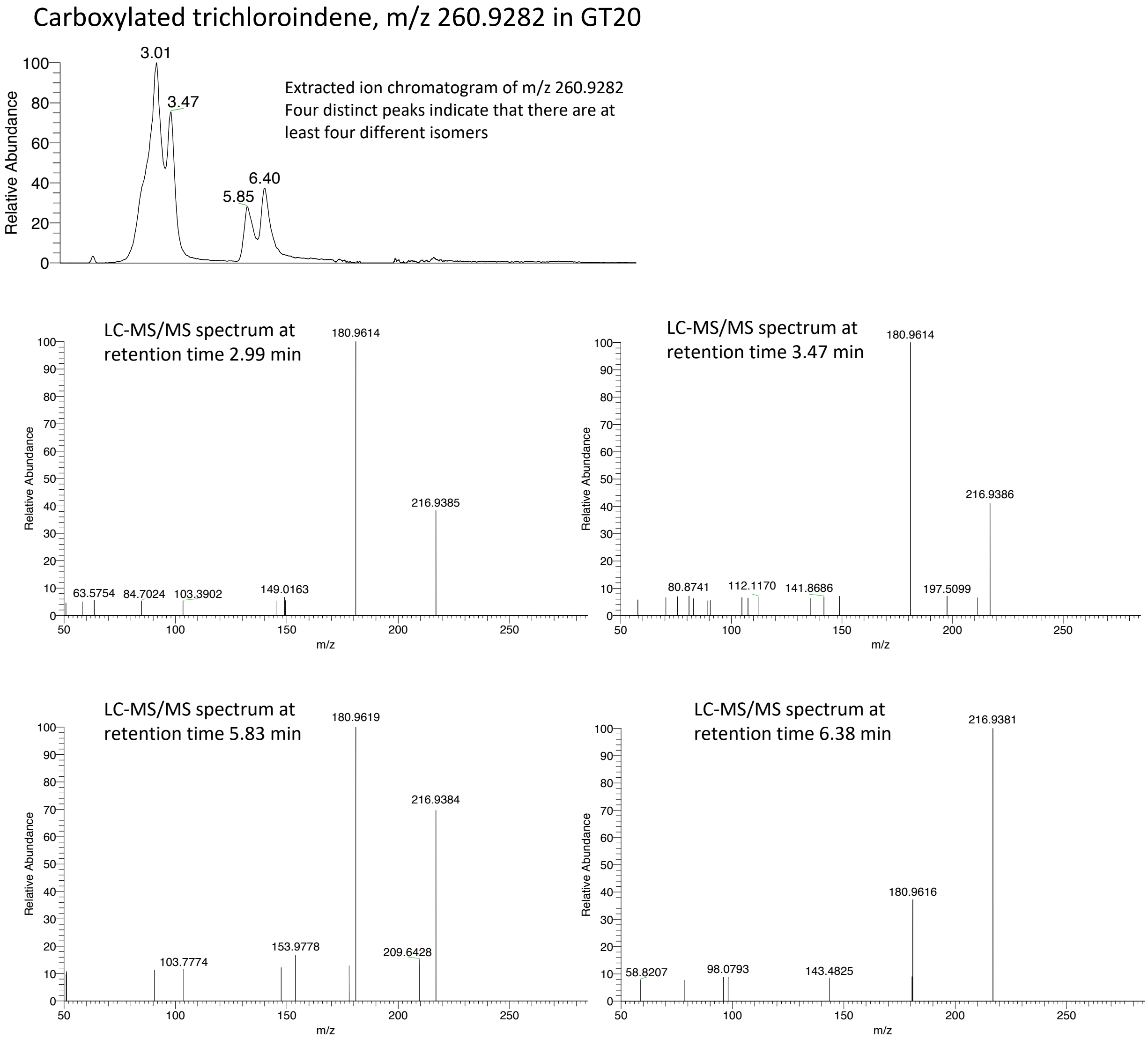


**Figure S4b**: Comparison of LC-MS/MS ion spectra of the fragmentation of m/z 260.9282 (carboxylated trichloroindene) at $t_R$ 2.99 min, 3.47 min, 5.83 min and 6.38 min (sample GT20).

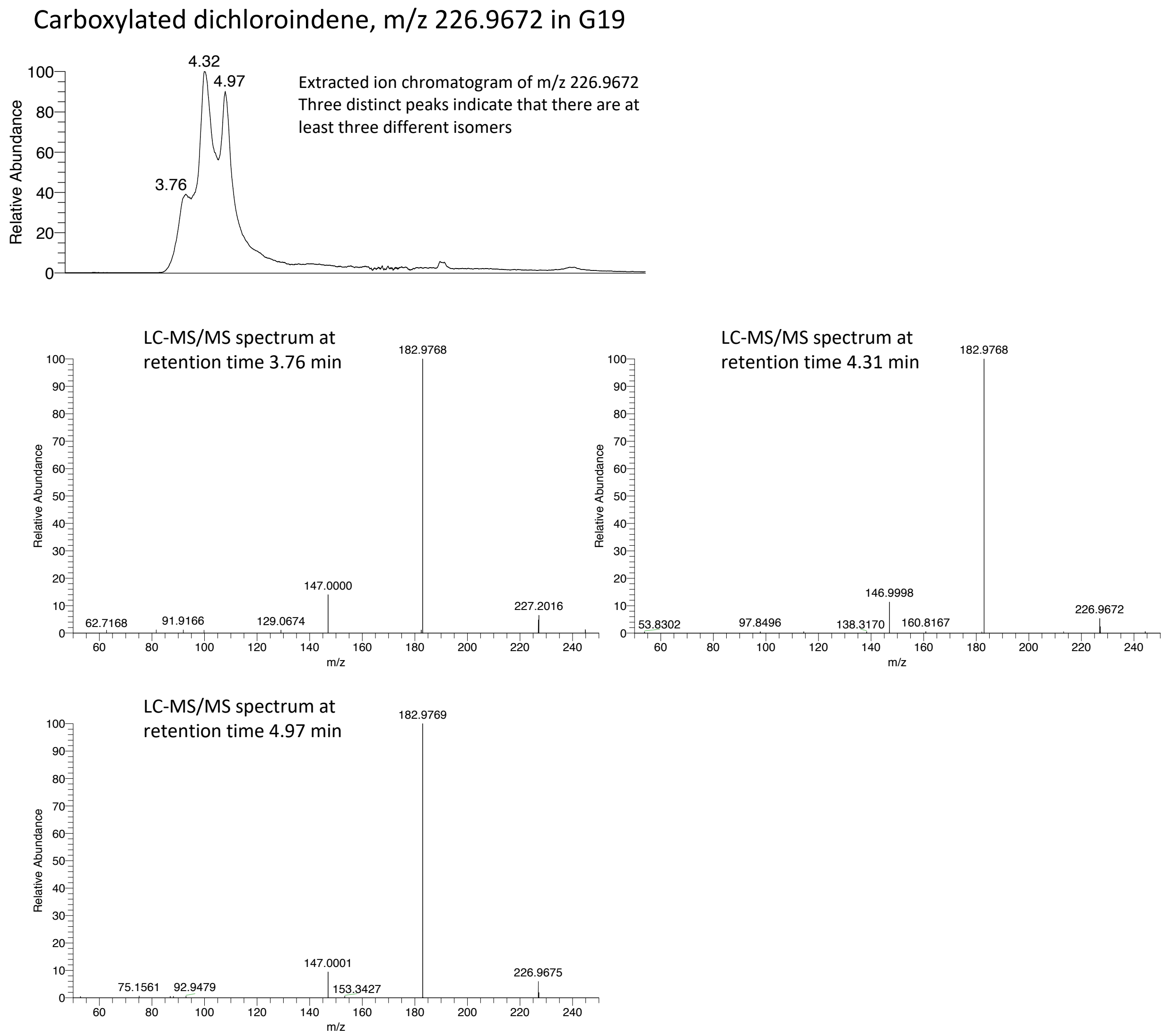


**Figure S4c**: Comparison of LC-MS/MS ion spectra of the fragmentation of m/z 226.9672 (carboxylated dichloroindene) at $t_R$ 3.76 min, 4.31 min and 4.97 min (sample G19).

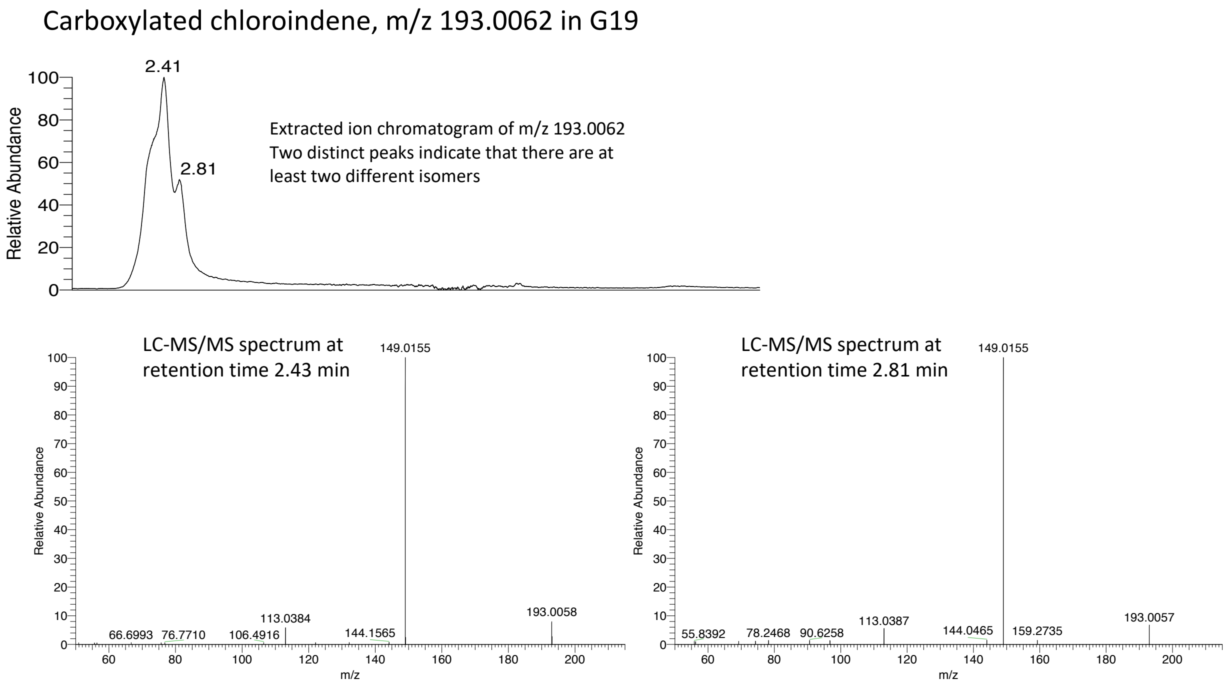


**Figure S4d**: Comparison of LC-MS/MS ion spectra of the fragmentation of m/z 193.0062 (carboxylated chloroindene) at $t_R$ 2.43 min and 2.81 min (sample G19).

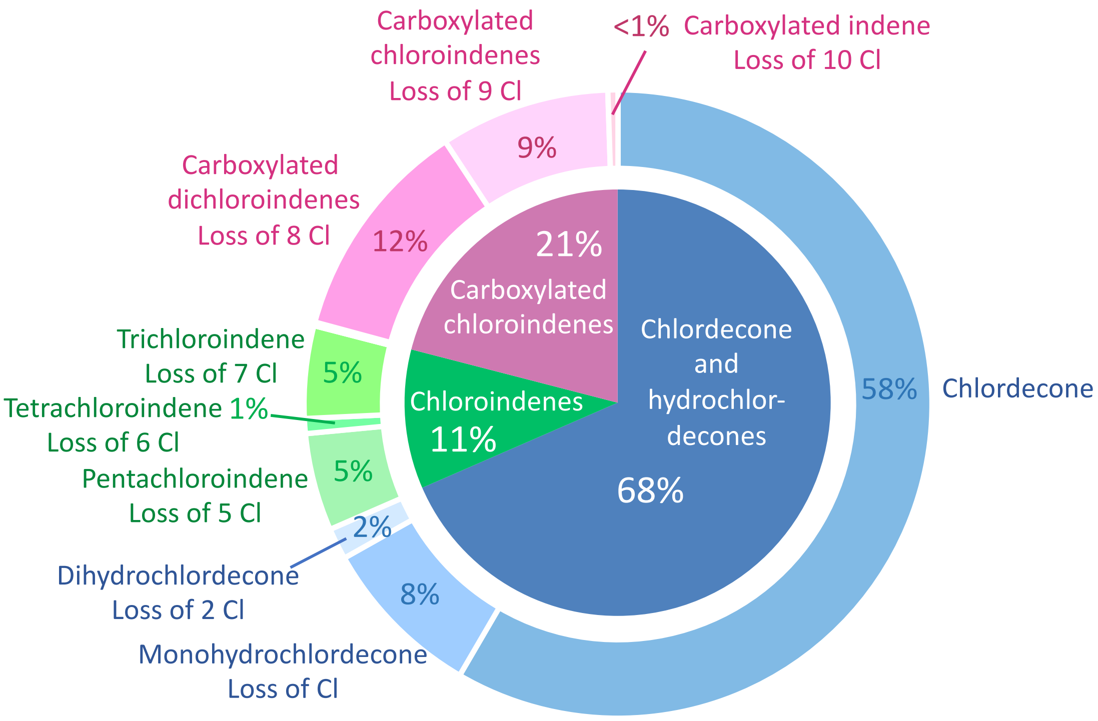


**Figure S5a:** Estimated distribution of chlordecone and dechlorinated metabolites in microcosm G4 (amended with chlordecone). Charts were constructed from measured and estimated concentrations in nmol/l (see Table S3b for details). Carboxylated trichloroindene was detected but with value less than 0.5%, and is not shown in the chart.

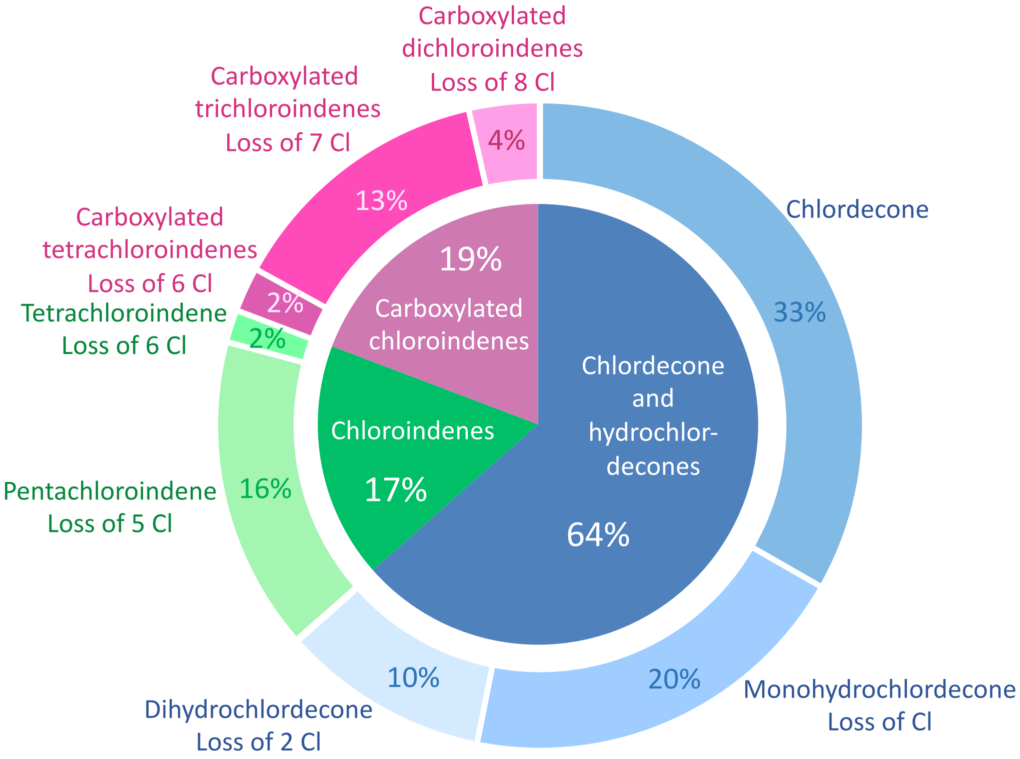


**Figure S5b:** Estimated distribution of chlordecone and dechlorinated metabolites in transfer culture GT3 (amended with chlordecone and TCE). Charts were constructed from measured and estimated concentrations in nmol/l (see Table S3b for details). Trihydrochlordecone and carboxylated chloroindene were detected but with values less than 0.5%, and are not shown in the chart.

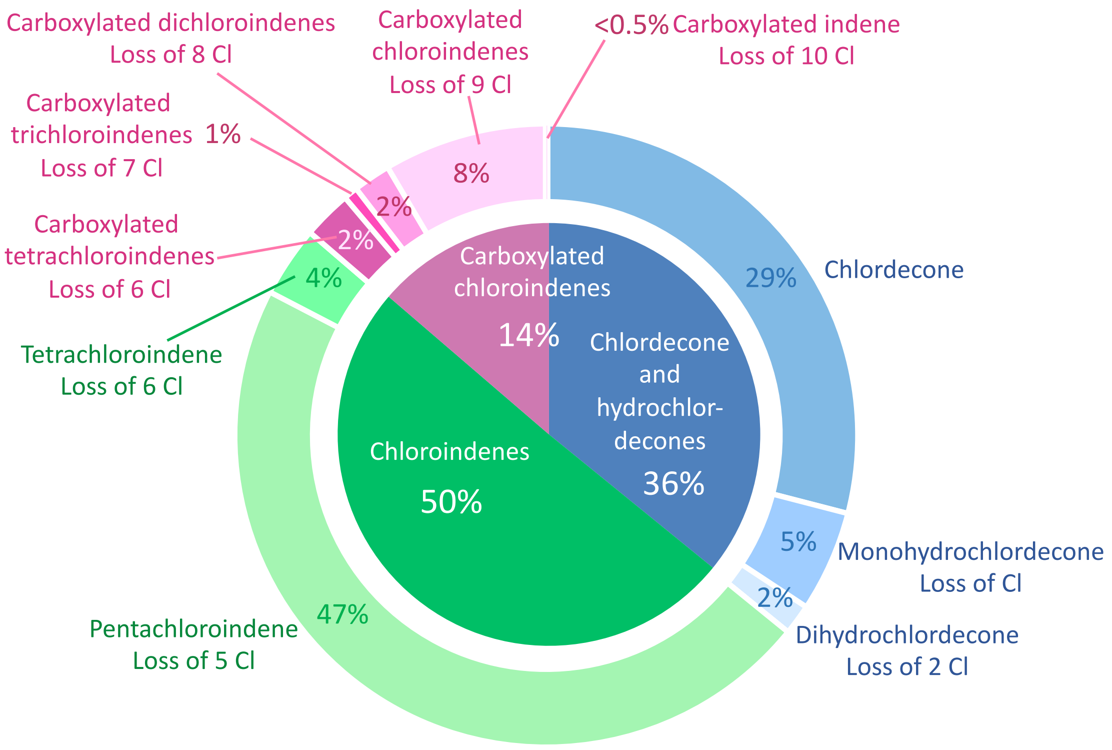


**Figure S5c:** Estimated distribution of chlordecone and dechlorinated metabolites in transfer culture GT5 (amended with chlordecone). Charts were constructed from measured and estimated concentrations in nmol/l (see Table S3b for details). Trihydrochlordecone was detected but with value less than 0.5%, and is not shown in the chart.

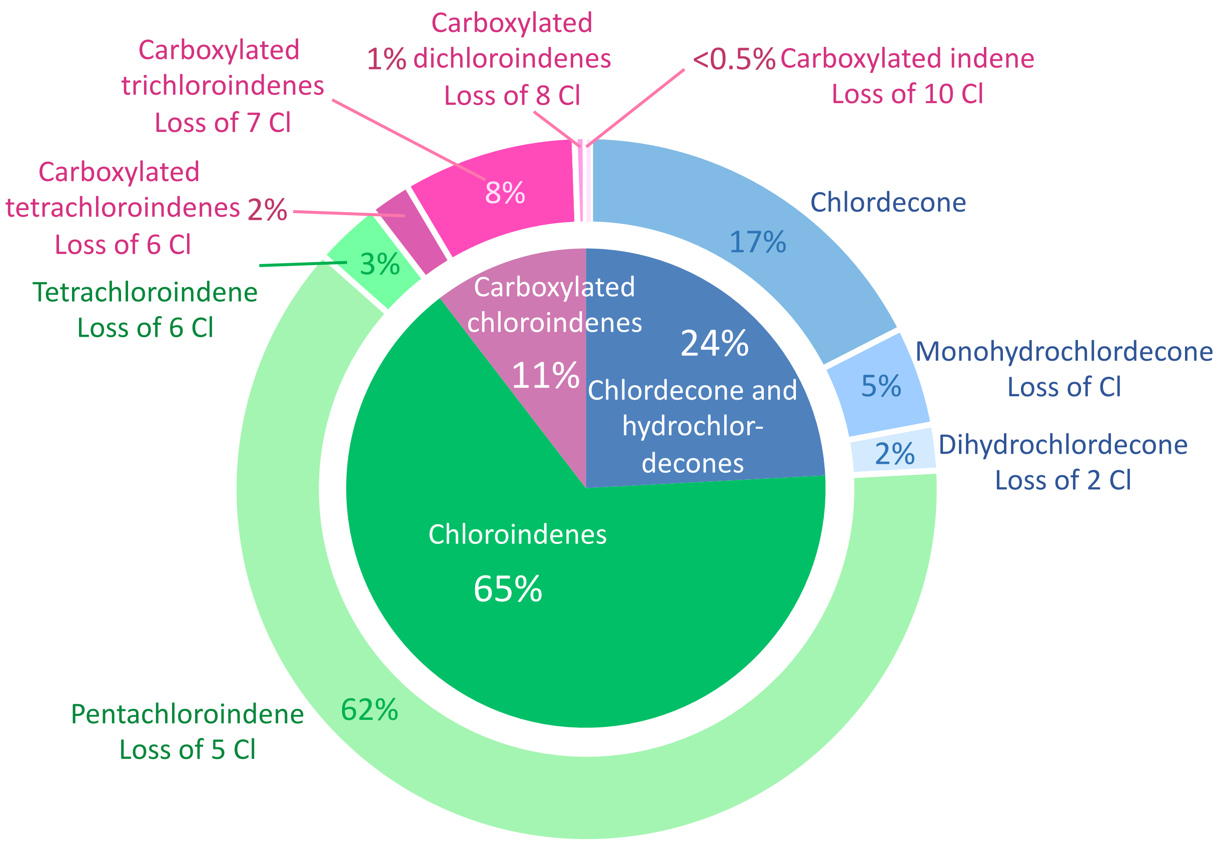


**Figure S5d:** Estimated distribution of chlordecone and dechlorinated metabolites in transfer culture GT33 (amended with chlordecone). Charts were constructed from measured and estimated concentrations in nmol/l (see Table S3b for details). Trihydrochlordecone and carboxylated chloroindene were detected but with values less than 0.5%, and are not shown in the chart.

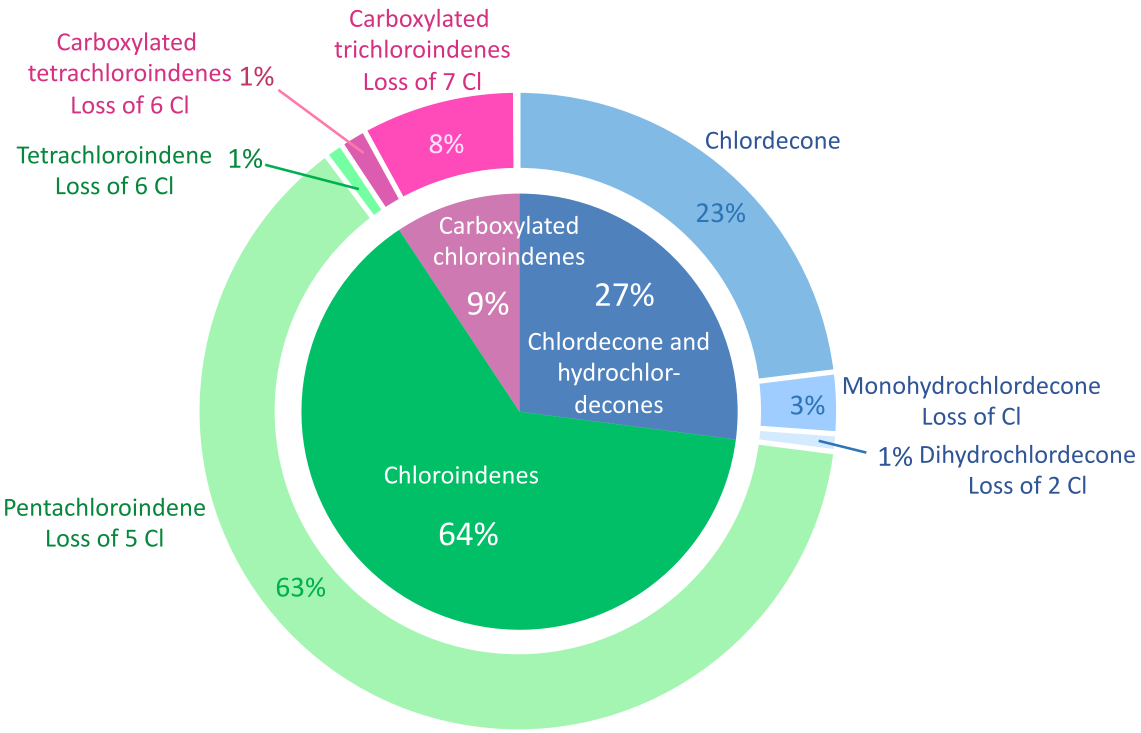


**Figure S5e:** Estimated distribution of chlordecone and dechlorinated metabolites in transfer culture GT4 (amended with chlordecone). Charts were constructed from measured and estimated concentrations in nmol/l (see Table S3b for details). Trihydrochlordecone, carboxylated dichloroindene and carboxylated chloroindene were detected but with values less than 0.5%, and are not shown in the chart.
